# Sex Determination in the garden lizard, *Calotes versicolor*: Is Environment a Factor?

**DOI:** 10.1101/312058

**Authors:** Priyanka, Rajiva Raman

## Abstract

The Indian garden lizard, *C. versicolor*, is known to lack sex chromosomes (Singh, 1974, Ganesh et al 1997). A report from the tropical southern India (Inamdar et al., 2012), claims a TSD (FMFM) mechanism in this species in which the male/female ratio in embryos oscillates within a range of 3-4°C. The present study presents results of experiments done in 4 consecutive breeding seasons of *C. versicolor* belonging to the subtropical/temperate climate of northern region of India. Eggs were grown at different temperatures (at 24.5/28/31.5°C) or under seminatural conditions. Another set of eggs was exposed to fadrozol, an aromatase inhibitor (AI), or Lithium Chloride (inhibitor of GSK3 enzyme inducing Wnt4-dependent Beta-Catenin). Results confirm our earlier finding that in this subtropical population of *C. versicolor*, temperature does not regulate gonadal differentiation but AI-induces sex reversal to the male sex (Ganesh and Raman, 1995, Ganesh et al 1999). We also report lack of any effect of Sox9 inhibitor on sexual differentiation which may be due to inadequate quantity or mode of application. Importantly, we report the serendipitous observation that in each year almost all the embryos/hatchlings were of the same sex ( females in 2013, ’15, ’16 and males in 2014) regardless of the rearing condition and duration of incubation. Obviously, parthenogenesis is not the cause of it. In the absence of an obvious reason to explain this pattern, we surmise that in this north Indian population of *C. versicolor*, female is the default sex, and certain epigenetic regulators could modulate the sexual differentiation of the individual.

## Introduction

*Calotes versicolor* is a common garden lizard distributed throughout south-east Asia, and prevalent in the Indian subcontinent. Male and females are easily distinguishable by virtue of the dorsal crest and hemipenis in the male, but chromosomally the two sexes are identical (Singh 1974, Ganesh et al. 1997). Also, the embryos grown at different temperatures do not indicate TSD (Ganesh and Raman, 1995). Thus, prima facie, *C. versicolor* lacks both CSD (Chromosomal Sex Determination) and TSD (Temperature Sex Determination), the two common modes of sex determination in reptiles, including lizards. However, treatment of growing eggs with androgens, the male gonad hormones, lead to all male progeny, especially with DHT, though the treatment with estradiol has no affect on the sex (Ganesh and Raman 1995, Ganesh et al 1999). Contrary to our results, a recent study on the same species from southern India shows a rather intricate TSD mechanism (Inamdar et al 2012). Using 14 different temperatures ranging from 23.5±0.5°C to35±0.5°C for egg incubation, the authors found all female progeny at low of 23.5 and high of 31.5±0.5^0^C and all male progeny at 25.5 and 34^0^C. The sex ratio was even at 24.5, 28.5 and 33^0^C, implying that between temperatures the sex ratio fluctuated through female-male-female-male (FMFM), and that there is TSD in this species.

We have re-examined sex ratios in the local population of *C. versicolor* under different incubation conditions including temperature. We did not find association of sex ratio with temperature, instead an unanticipated set of results in the delivery of single sex progeny in successive breeding seasons were observed. We present the results in this paper.

## Materials & Methods

*C. versicolor* is dispersed throughout India. It lives in small bushes even in domestic lawns, and feeds on small insects. Not being an endangered species, there are no specific restrictions on their collection from the wild. It is a seasonal breeder with prolific reproductive activity during monsoon. However, they are not easily amenable to captive breeding. Laboratory-born hatchlings rarely survive beyond 15 days. Varanasi, where the study was performed, is situated in the eastern zone of north India (latitude north: 25° 8’; longitude east: 83° 1’), and experiences subtropical climate with reasonably cold winters and strong summers, intervened by monsoon rains between the end of June and mid-September which induces reproductive activity in the lizard. Therefore each year during this period, a fresh lot of animals were collected from the wild (forests around 20 kms away from the University campus), and brought to the lab. All applicable international, national and/or institutional guidelines for the care and use of animal were followed. Following 2-3 days of acclimatization in wooden cages, 10-12 gravid females were guillotined for collection of eggs (a clutch 12-20 eggs per individual) and other tissues. The eggs were placed in earthen pots having sterile sand, and - barring one group that was placed in a bush ~200 meters away from the lab - all others were maintained in BOD incubators at different temperatures (28^0^C/31.5^0^C/24.5^0^C all ±0.5^0^C). Fadrozole (Aromatase Inhibitor AI; 5μg/egg) or Lithium Chloride (50mMol LiCl, 5μl/egg), a GSK3 inhibitor that activates β Catenin resulting in silencing of *Sox9* (Bernard et al., 2012), were topically applied to certain eggs on day 6 of incubation. Water was sprinkled every morning and evening in the pots, to maintain humidity. Diurnal weather charts of the University campus were obtained from the Meteriological Department in the University. Mesonephros Gonadal complex (MGC) of certain embryos and hatchlings were fixed in Bouin and preserved for gonadal histology. Tissue sections were stained with haematoxylene.

Since embryonic gonads in this species, retained the testis and ovarian remnants until hatching, embryonic sex was deciphered only late in development through presence (Hp+) or absence (Hp^−^) of hemipenis (Hp) that emerges in all the embryos by about day 30 but regresses in putative females latest by day 40.

## Results and Discussion

The study was done in 4 consecutive breeding seasons (2013 - 2016), and different types of experiments were set up in different years with the objective of finding the sex of individuals grown under those conditions (temperature, quasi-natural, and gonadal enzyme inhibitors). We present the results year-wise, as they were obtained, because each year guided the experiments for the following year (see Table 1).

**Table 1.** Year-wise details of the embryos/hatchlings with and without hemipenis (Hp) under different experimental conditions

| Year | Incu. Temp | Treatment | Total eggs | Eggs Survived | Day sacrificed | Hp<sup>+</sup> | Hp<sup>-</sup> |
| --- | --- | --- | --- | --- | --- | --- | --- |
| 2013 | 28<sup>o</sup>c | Fadrozol (5ug/egg) | 17 | 16 | 42 | 14 | 2 |
|  |  | Control | 11 | 11 | 42 | 00 | 11 |
|  | 28<sup>o</sup>c | LiCl (50mM; 5ul/egg) | 15 | 14 | 41 | 01 | 13 |
|  |  | Control | 16 | 16 | 41 | 00 | 16 |
|  | 31.5<sup>o</sup>c | High temperature | 15 | 15 | 40 | 1 | 14 |
| 2014 | 28<sup>o</sup>C | Fadrozol (5ug/egg) | 10 | 08 | 42 | 08 | 00 |
|  |  | Control | 10 | 09 | 42 | 09 | 00 |
|  | 28<sup>o</sup>C | LiCl (50mM; 5ul/egg) | 11 | 11 | 41 | 11 | 00 |
|  |  | Control | 13 | 13 | 41 | 13 | 00 |
| 2015 | ambient | In a bush in nature | 35 | 31 | 43 | 00 | 10 |
|  |  |  |  |  | 44 | 00 | 04 |
|  |  |  |  |  | hatchling | 00 | 13 |
|  | 31.5<sup>o</sup>C | High temperature | 10 | 10 | hatchling | 00 | 10 |
|  | 28<sup>o</sup>C | Control | 10 | 09 | 43 | 00 | 09 |
|  |  |  | 07 | 07 | hatchling | 00 | 07 |
|  | 24.5<sup>o</sup>C | Low temperature | 14 | 10 | 66 | 07* | 00 |
|  |  |  |  |  | 80 | 00 | 02 |
|  |  |  |  |  | hatchling | 00 | 01 |
| 2016 | 28<sup>o</sup>C | Control | 30 | 30 | 37, 39,<br>41, 44,<br>47 (06<br>each day) | 00 | 30 |
|  | 31.5<sup>o</sup>C | High Temperature | 15 | 12 | 40<br>Hatchling | 00<br>05 | 04<br>03 |

\* These embryos were small and equivalent to those at days 30-35 at 28<sup>o</sup>C, the stage at which all the embryos have rudimentary Hemipenis. Presence of Hp at this stage does not ensure

In the year 2013, the eggs were incubated at 31.5°C (expected 100%♀) and 28^0^C (♂/♀ 50% each; standard temperature). In addition, certain eggs were topically applied fadrozol (Aromatase Inhibitor, AI) or LiCl (Lithium chloride) on day 6 ( at 28^0^C). Each group comprised at least 10 eggs. Embryos were sacrificed mostly between days 41 and 43 of development, and checked for presence or absence of hemipenis (Hp).

The expectation was that AI-treated eggs would be all males, those treated with LiCl or kept at 31.5°C would be all females, and the male:female ratio in 28^0^C embryos will be even.

The results show (Table 1) that while ~90% of the AI-treated eggs were Hp^+^, the LiCl and the 31.5°C embryos were all Hp^−^. This result was on the expected line except for the fact that the 28^0^C (control) embryos also were all Hp^−^ when they were expected to be a mix of Hp^+^ and Hp^−^ embryos. Thus barring the AI treated embryos (14♂:2♀), most others were females (53♀:2♂). We repeated the AI and LiCl-treatments in the next breeding season (year 2014). This year, while the AI-treated eggs were all Hp^+^ (8♂:0♀), those treated with LiCl and the controls (28^0^C) too were Hp^+^ (33♂:0♀), unlike in 2013 when they were females. Clearly, AI-treatment led to reversal of sex towards male in both the years as shown earlier (Ganesh et al 1999). In contrast, LiCl results were identical to the controls in both the years. Apparently LiCl did not affect gonadal differentiation, possibly either because of the embryonic stage at which it was administered or due to the mode of application or else because of inadequate concentration. Thus in both the years barring the AI-treated, almost all the embryos were of one sex, regardless of the environment. However, two precociously gravid females caught in the first week of June (ambient temp. ~45^0^C, when gravid females are rarely seen), and had just 7 and 5 eggs, yielded 4 Hp^+^ and 8 Hp^−^ embryos.

In the next breeding season (year 2015), embryos were incubated at 24.5°C or 31.5^0^C. Another bunch of eggs was placed in a bush some 200 meters away from the lab, in earthen pots in sterile sand, covered with a fine net (quasi-natural condition). They were thus exposed to diurnal fluctuations of temperature (7-10^0^C - between 37 and 22^0^C), humidity, day-night length etc. In another variation to the protocol, only 14 bush grown and 10 controls were sacrificed as embryos on days 43 & 44, rest were allowed to grow full term to hatch so that the sex could be confirmed unambiguously in the hatchlings by examining the Hp and the gonadal histology. The bush-grown embryos hatched between days 47-48 while those reared at 31.5^0^C hatched on days 55-56. Each embryo and hatchling was Hp^−^. Gonadal sections of hatchlings or embryos observed under microscope displayed growing ova and no sex cords, confirming their ovarian nature (Fig. 1). The 24.5^0^C embryos did not hatch until much later. Therefore we dissected 7 of them on day 66, which developmentally compared to the 30-35 day old 28^0^C embryos. They, as expected of embryos at that stage, had rudimentary Hps. The 2 embryos dissected on day 80 (comparable to day 40 embryo at 28^0^C), and the one that hatched on day 86 were rather weak but all of them were lacked Hp. Considering that all other embryos and hatchlings were females, we inferred that the 24.5^0^C-day 66-Hp^+^ embryos to be females too. But even if we exclude them as ambiguous, there were 56 females against 00 males, irrespective of the environment they grew. The same was the condition in 2016 where embryos were grown at 28 or 31.5^0^C. Those kept at 28^0^C were dissected serially on days 37, 39, 41, 44 and 47 (6 each n= 30) with the objective of catching the period when rudimentary Hemipenis that emerged around day 30 of development atrophied in prospective female embryos, but all of them, even those of day 37, turned out to be Hp^−^. Of the 31.5^0^C lot, all 4 the embryos were Hp^−^, but of the 5 hatchlings were Hp^−^ as against 3 Hp^+^ hatchlings.

**Fig. 1.**
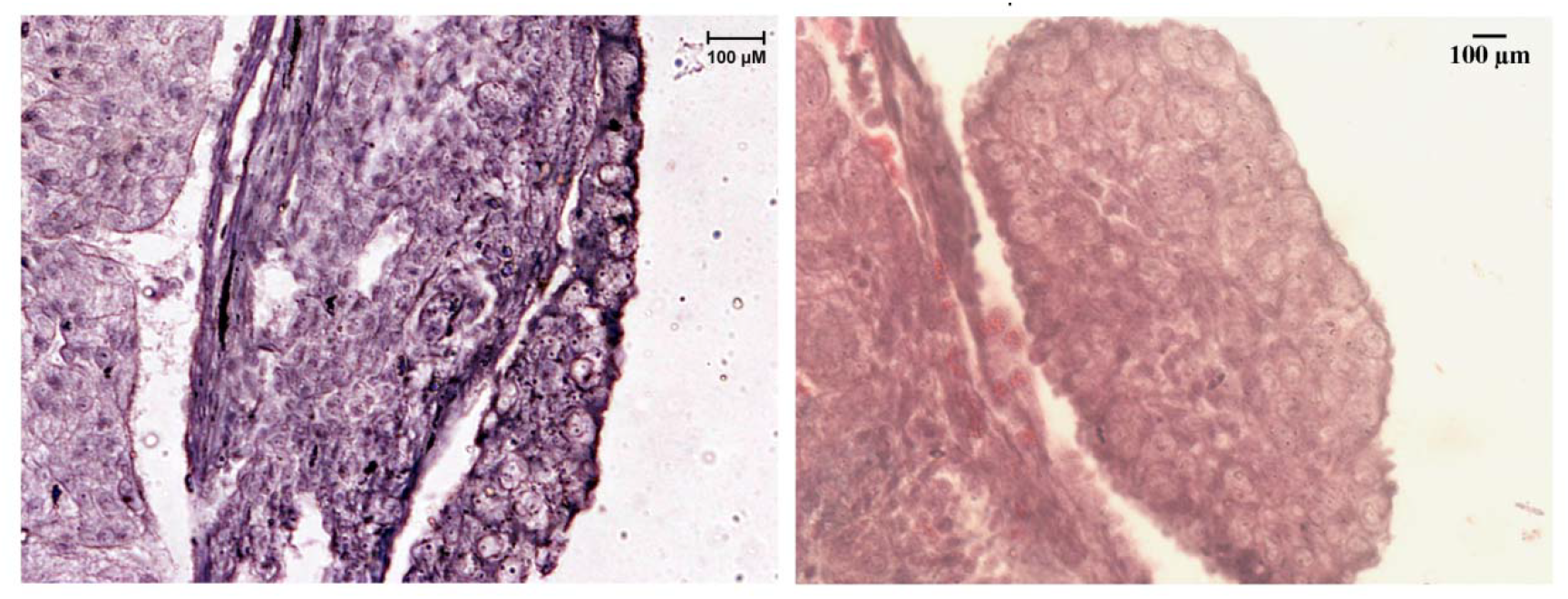
Hematoxylene-stained 7μm thick sections of the gonads of (a) 10 day old hatchling of embryo grown at 28°C and (b) 44 day old embryo grown in natural condition. Well developed cortex with large meiocytes is seen in both sections. While there is no medullary remnant in the embryonic gonad (b), atrophied medulla with no sex cords is seen in the hatchling (a). 1. Hematoxylene-stained 7μm thick sections of the gonads of (a) 10 day old hatchling of embryo grown at 28°C and (b) 44 day old embryo grown in natural condition. Well developed cortex with large meiocytes is seen in both sections. While there is no medullary remnant in the embryonic gonad (b), atrophied medulla with no sex cords is seen in the hatchling (a).

Data on the diurnal climatic conditions in and around the University campus during these 4 ywars were obtained from the Materiological Department. As expected, June was the hottest and the driest month, up to the third week, in all the 4 years. A comparison of the mean of minimum, maximum and the diurnal range of temperatures and relative humidity between June and August, showed no difference in the 4 years except that the rain fall was the lowest in 2014 (593mm) and maximum in 2016 (962mm). There were only 27 days of rain in 2014, the year in which all the embryos were males (Table 2).

**Table 2.** A chart summarising diurnal weather conditions through June and August during the 4 years of study

| S. No | Year | Min. Temp. Range (Av.) | Max. Temp. Range (Av.) | Fluctuation diurnal temp. | % Relative Humidity | Range Rain-fall (mm) | Days of rainfall |
| --- | --- | --- | --- | --- | --- | --- | --- |
| 1 | 2013 | 24.4-28.6<br>(26.3) | 29.8-39<br>(34.1) | 7-10<sup>0</sup> C | 54-100 | 681.5 | 44/92 |
| 2 | 2014 | 22.9-27.9<br>(25.7) | 29.6-39<br>(34.1) |  | 54-100 | 573 | 27/92 |
| 3 | 2015 | 25-28.7<br>(26.8) | 27.2-38<br>(33.8) |  | 67-100 | 768 | 33/92 |
| 4 | 2016 | 24.2-31.9<br>(27.4) | 28.2-43<br>(35.0) |  | 32-100 | 963.3 | 47/92 |

Thus the present set of experiments done over a period of 4 years, exposing eggs to different regimes of constant temperatures as well as quasi-natural conditions in which environment fluctuated diurnally, give no reason to suspect Temperature or Environment dependent sex determination (TSD/ESD) mechanism in this population of *C. versicolor*. Therefore the only way to reconcile the present results with that of Inamdar et al (2012) would be the different climatic conditions the two populations live in; the prolonged breeding season and no exposure to extreme climatic in tropical environs of southern India differing from their north Indian morph. The present results however have added complexity insofar that all the progeny are of the same sex in consecutive 4 years. It is particularly strange because we have not recorded similarly skewed sex ratio in the previous 15 years. Independent of us, there is at least one report on the sex ratio of the lab-born hatchlings of this species from this region for the calendar years 1964-1966 which recorded 159 males to 100 females (Pandha and Thapliyal, 1967).

Naturally occurring skewed sex ratios and sex reversals are not uncommon in animal kingdom, more specifically in reptiles. In fact they are considered a source of ongoing evolution of sex determining mechanisms (Holleley et al., 2015). However, birth of only one sex individuals in nature is rare except due to parthenogenesis (Maslin, 1971). In sexually reproducing species, parthenogenesis is noticed in populations recording absence of males, but that seems improbable in this species because of the natural cohabitation of males and females in this population, and also because of birth of all male progeny in 2014. Several other reasons/mechanisms of skewed sex ratio in reptilian populations have been shown, largely drawn from the skewed ratio in adult populations (Charnov, 1982; Trivers & Willard 1973; West et al., 2002). There are also examples of prefertilisation adaptation towards the sex ratio of the offspring depending upon the accumulation of the gonadal hormones in yolk during maturation of egg (Radder, 2007). Besides environmental factors, examples of the lizards, *Ctenophorus pictus* and *Anolis sagrei* allude to a genetic mechanisms responsible for skewed sex ratio (Olsson et al., 2007, Cox & Calsbeek, 2010)..

Previous studies from our lab had shown lack of TSD in *C. versicolor*, but they also showed androgen-induced unilateral sex reversal towards male progeny (Ganesh and Raman 1995). Therefore, we had surmised that sex determination in *C. versicolor* is under weak genetic regulation which could be superseded by hormonal interventions. We also proposed that female is the default sex in *C. versicolor* whose fate is fixed later than that of male during development, and if the male pathway genes are switched on or induced (say, by androgens) earlier than the fixation of the female pathway, the individuals would develop into males (Ganesh et al 1999, Chakraborty and Raman 2010). Even though at present there is no obvious and cogent explanation to the unisex progeny reported here, it does lead to a possibility that the reason for the default nature of the female sex could be due to accumulation of gonadal hormones in the egg yolk or imprinting of certain loci in the gametes which could control the sex ratio of the embryo, possibly in response to population dynamics and/or prevailing environment. We record this novel observation in this widely distributed species so that future investigations on this species take into account occasional extreme skewing of sex ratios.

## Acknowledgements

This work was supported by grants from the Department of Science & Technology and later by the Council of Scientific & Industrial Research, New Delhi under the Scientist Emeritus scheme no. 21(0946)/13/EMR-II to RR. Priyanka thanks CSIR for her Junior and Senior Research Fellowships.

## References

Bernard P, Ryan J, Sim H,. Czech DP, Sinclair AH, Koopman P, Harley VR (2012) Wnt signaling in ovarian development inhibits Sf1 activation of Sox9 via Tesco Enhancer. Endocrinology 153:901–912.

Chakraborty A. Raman R (2010) Modulation of gene activity in androgen-induced sex reversal in the garden lizard, *Calotes versicolor*. Sexual Dev 4, 162–169.

Charnov EL(1982) The theory of sex allocation. Princeton University Press, Princeton.

Cox RM, Calsbeek R (2010) Cryptic sex-ratio bias provides indirect genetic benefits despite sexual conflict. Science 328:92–94.

Ganesh S, Raman R (1995). Sex reversal by testosterone and not by estradiol or temperature in *Calotes versicolor*; the lizard lacking sex chromosomes. Journal of Experimental Zoology 271:139–144.

Ganesh S, Mohanty J, Raman R (1997) Male biased distribution of the human Y chromosomal genes SRY and ZFY in the lizard, *Calotes versicolor*, which lacks sex chromosomes and temperature dependent sex determination. Chrom Res 5: 413–419.

Ganesh S, Choudhary B, Raman R (1999). Temporal difference between testis and ovary determinations with possible involvement of testosterone and aromatase in gonadal differentiation in TSD lacking lizard, *Calotes versicolor*. J Exp Zool 283:600–607.

Holleley C. E., O’Meally D, Sarre SD, Graves JAM, Ezaz T, Matsubara K, Azad B, Zhang X, and Georges A (2015) Sex reversal triggers the rapid transition from genetic to temperature-dependent sex. Nature 523:79–81.

Inamdar LS, Vani V, Seshagiri PB (2012) A tropical oviparous lizard, *Calotes versicolor*, exhibiting a potentially novel FMFM pattern of temperature dependent sex determination. J Exp Zool 317:32–46.

Maslin TP (1971) Parthenogenesis in reptiles. American Zoologist 11:361–380.

Olsson M, Shine R (2001) Facultative sex allocation in snow skink lizards (Niveoscincus microlepidotus). J Evol Biol 14:120–128.

Pandha SK, Thapliyal JP (1967) Egg laying and development in the garden lizard, *Calotes versicolor*. Copiea 1967:121–125.

Radder RS (2007) Maternally derived egg yolk steroid hormones and sex determination: Review of a paradox. J Biosci 32:1213–1220.

Singh L (1974) Study of mitotic and meiotic chromosomes in seven species of lizards. Proc Zool Soc Calcutta 27:57–79.

Trivers RL, Willard DE (1973) Natural selection of parental ability to vary the sex ratio of offspring. Science 179:90–92.

West SA, Reece SE, Sheldon BC (2002) Sex ratios, Heredity 88:117–124.

